# Functional genetic validation of key genes conferring insecticide resistance in the major African malaria vector, *Anopheles gambiae*

**DOI:** 10.1101/749457

**Authors:** Adriana Adolfi, Beth Poulton, Amalia Anthousi, Stephanie Macilwee, Hilary Ranson, Gareth J Lycett

**Affiliations:** Vector Biology Department, Liverpool School of Tropical Medicine, Liverpool L3 5QA, UK

**Keywords:** Insecticide resistance, Cytochromes P450, Glutathione-S-transferase, GAL4/UAS, *Anopheles*, Functional analysis

## Abstract

Resistance in *Anopheles gambiae* to members of all four major classes (pyrethroids, carbamates, organochlorines and organophosphates) of public health insecticides limits effective control of malaria transmission in Africa. Increased expression of detoxifying enzymes has been associated with resistance, but direct functional validation in *An. gambiae* has been lacking. Here we perform transgenic analysis using the GAL4/UAS system to examine insecticide resistance phenotypes conferred by increased expression of the three genes - *Cyp6m2, Cyp6p3* and *Gste2 -* most often found upregulated in resistant *An. gambiae*. We report the first evidence in *An. gambiae* that organophosphate and organochlorine resistance is conferred by overexpression of GSTE2 in a broad tissue profile. Pyrethroid and carbamate resistance is bestowed by similar *Cyp6p3* overexpression, and *Cyp6m2* confers only pyrethroid resistance when overexpressed in the same tissues. Conversely, such *Cyp6m2* overexpression increases susceptibility to the organophosphate malathion, presumably due to conversion to a more toxic metabolite. No resistant phenotypes are conferred when either *Cyp6* gene overexpression is restricted to the midgut or oenocytes, answering long standing questions related to the importance of these tissues in resistance to contact insecticides. Validation of genes conferring resistance provides markers to guide control strategies, and the observed negative cross-resistance due to *Cyp6m2* gives credence to proposed dual insecticide strategies to overcome pyrethroid resistance. These trasnsgenic *An. gambiae* resistant lines are being used to test potential liabilities in new active compounds early in development.

**SIGNIFICANCE STATEMENT:** Insecticide resistance in *Anopheles gambiae* mosquitoes can derail malaria control programs, and to overcome it we need to discover the underlying molecular basis. Here, for the first time, we characterise three genes most often associated with insecticide resistance directly by their overproduction in genetically modified *An. gambiae*. We show that overexpression of each gene confers resistance to representatives of at least one insecticide class and, taken together, the three genes provide cross-resistance to all four major insecticide classes currently used in public health. These data validate the candidate genes as markers to monitor the spread of resistance in mosquito populations. The modified mosquitoes produced are also valuable tools to pre-screen new insecticides for potential liabilities to existing resistance mechanisms.

## INTRODUCTION

From the year 2000 until recently, the number of worldwide malaria cases had steadily fallen mainly due to the widespread rollout of insecticide treated bed nets in endemic areas (1, 2), which offer protection against bites from *Plasmodium* infected *Anopheles* mosquitoes. There is growing evidence suggesting that the stalling in malaria control can be at least partially attributed to the increasing levels of insecticide resistance in *Anopheles* vectors (3). Resistance in dominant African *Anopheles* vectors has been recorded to all major insecticide classes currently used in public health (pyrethroids, organochlorines, carbamates and organophosphates) (4). Therefore, understanding the mechanisms by which mosquitoes evolve resistance is critical for the design of mitigation strategies and in the evaluation of new classes of insecticides.

Research into the molecular mechanisms that give rise to resistance in mosquitoes have identified target site modifications and increased metabolic detoxification as the two main evolutionary adaptions (5), that often co-exist in *An. gambiae*. Families of detoxification enzymes, including cytochromes P450 (CYP) and glutathione-S-transferases (GST), can provide phase I metabolism of insecticides and phase II conjugation reactions that alter the toxicity of compounds and increase polarity, enhancing excretion (6, 7).

To identify and characterise the role of the causative resistance genes from these detox families, a sequential process of transcriptomic, proteomic and *in vivo* functional analysis is often applied (8). Candidate genes with upregulated transcription or strong signatures of selection in resistant mosquitoes are typically expressed in bacteria to provide evidence of insecticide depletion and/or metabolism *in vitro* (9–19). Further studies have used the *Drosophila* transgenic model to determine whether expression of single *Anopheles* genes confers increased tolerance to insecticides (13–18, 20).

This workflow has implicated a role in resistance of two cytochrome P450 genes, *Cyp6m2* and *Cyp6p3*, and a Glutathione S Transferase gene, *Gste2*, that are consistently upregulated in resistant field populations found across Africa (21). However, there are often discrepancies in results from recombinant protein activity and transgenic *Drosophila* analyses. For example, while expression studies of *Cyp6m2* and *Cyp6p3* in *E. coli* (10, 11) and *Drosophila* (15) suggest that both gene products can detoxify pyrethroids, the two systems produce conflicting results in respect to carbamate (15) and organochlorine insecticide detoxification (12, 15, 19). Moreover, the involvement of *An. gambiae and An. funestus Gste2* orthologues in resistance to pyrethroid insecticides has produced contradictory results when explored in *Drosophila* (16, 20).

Clearly, functional validation of *Anopheles* genes directly in the mosquito would provide the benchmark approach to address these questions, however to date transgenic tools to perform such analysis have been limited. To this end, we have developed the GAL4/UAS expression system in *An. gambiae* (22–24) which allows genes to be overexpressed in a susceptible mosquito background and for resultant resistance phenotypes to be examined using the standard insecticide assays that have been developed for comparative analysis in mosquitoes by WHO (25).

*In vivo* functional analysis in *Anopheles* can also help discover the mosquito tissues that are specifically involved in insecticide metabolism. Our previous research indicated high P450 activity in the midgut and oenocytes, since the essential P450 co-enzyme CPR is highly expressed in these tissues, and RNAi knockdown of *Cpr* increased mosquito sensitivity to a pyrethroid insecticide (26). Moreover, *Cyp6m2* has been reported as enriched in the *An. gambiae* midgut (11) and *Cyp6p3* was found upregulated in midguts from pyrethroid resistant populations (27).

Here we have used the GAL4/UAS system to overexpress *Cyp6m2* or *Cyp6p3* genes in multiple tissues or specifically in the midgut or oenocytes of a susceptible *An. gambiae* strain and assayed the modified mosquitoes against representatives of each insecticide class available for public heath use. In doing so, we determined the resistance profile generated for each gene and compared these results to those obtained in *Drosophila* and *in vitro*. We then analysed the other major candidate, *Gste2*, to examine its role in conferring DDT resistance and also extending its testing to other classes of insecticides in which its role has yet to be tested *in vivo*.

In this work, we report the first use of the GAL4/UAS system in *Anopheles* as a benchmark to determine whether single candidate genes and/or expression in individual tissues are able to confer WHO-defined levels of resistance to the four public health classes of insecticides, including for the first time organophosphates. Crucially we find that, when assayed in *An. gambiae*, overexpression of *Cyp6m2, Cyp6p3* or *Gste2* produce cross-resistance phenotypes that encompass members of all four classes of insecticides currently used for malaria control.

## RESULTS

### Mosquito lines generated for UAS-regulated expression of Cyp6m2 and Cyp6p3

YFP marked UAS-*Cyp6m2* and -*Cyp6p3* lines were created by site directed recombination mediated cassette exchange (RMCE) into a docking (CFP:2x*attP*) line A11 (24) to produce mosquitoes carrying transgene insertions in the same genomic site. By mitigating for genomic position effects, this allows more reliable comparison of the effects of *Cyp6m2* and *Cyp6p3* overexpression on resistance.

A summary of the screening and crossing strategy used to create the UAS responder lines is illustrated in Table 1. RMCE results in canonical cassette exchange in two potential orientations, however integration of the whole donor transgene can also occur into either *attP* site. Fluorescent marker screening of F_1_ progenies from F_0_ pooled mosquitoes revealed that cassette exchange and integration events occurred in all experiments as shown by the recovery of individuals carrying single (YFP: exchange) or double (CFP/YFP: integration) markers (Table 1).

**Table 1.** Summary of the screening and crossing strategy adopted to create and establish the UAS responder lines by RMCE.

| Docking line<br>(No. Embryos) | F<sub>0</sub> pools<br>(No. and sex) | F<sub>0</sub><br>isofemale | F<sub>1</sub> transgenics |  | Orientation of cassette<br>exchange** |
| --- | --- | --- | --- | --- | --- |
|  |  |  | YFP+ | YFP+/CFP+ |  |
| **A11_UAS-**<br>***Cyp6m2***<br>(347) | M2-1<br>(24 ♀) | G | 0 | 2 | N/A |
|  |  | J | 2♂ | 0 | 2 F<sub>1</sub> ♂ - A |
|  | M2-2<br>(25 ♂) | N/A | 0 | 0 | N/A |
| **A11_UAS-**<br>***Cyp6p3***<br>(460) | P3-1<br>(28 ♀) | N/A | 7♀, 4♂ | 1 | 5 F<sub>1</sub> ♀ - A x2, B x3 |
|  | P3-2<br>(27 ♀) | N/A | 2♀, 8♂ | 2 | 2 F<sub>1</sub> ♀ - A, B |
|  | P3-3<br>(13 ♀) | N/A | 0 | 0 | N/A |
|  | P3-4<br>(56 ♂) | N/A | 10♀, 13♂ | 4 | 3 F<sub>1</sub> ♀ - A, B x2 |
| **Ubi-A10_**<br>**UAS-*Gste2***<br>(208) | E2-1<br>(10 ♂) | N/A | 0 | 0 | N/A |
|  | E2-2<br>(12 ♀) | N/A | 0 | 0 | N/A |
|  | E2-3<br>(19 ♂) | N/A | 2♂ | 36♀, 44 ♂ | 2 F<sub>1</sub> ♂ - A |
|  | E2-4<br>(24 ♀) | A | 3♀, 3♂ | (7)* | F<sub>2</sub> progeny of 1 F<sub>1</sub> ♂ - B |
|  |  | E | 4♀, 3 ♂ | 2♀, 2♂ | 1 F<sub>1</sub> ♀ - A<br>F<sub>2</sub> progeny of 1 F<sub>1</sub> ♂ - A |

\*did not survive to adulthood.
\*\*As cassette exchange may occur in two different orientations with respect to the chromosome, designated A or B, orientation check was performed on F<sub>1</sub> YFP-positive individuals or on the F<sub>2</sub> progeny deriving from single YFP-positive individuals.

Molecular analysis revealed one exchange orientation (A) in transgenic UAS-m2 individuals and both orientations for UAS-p3 transformation as indicated by diagnostic PCR (Fig. S1). Overall, we found at least two events for UAS-m2 transformation, having equal efficiencies of 2% for cassette-exchange and integration (1/49 F_0_ founders); while for the UAS-p3 transformation, at least nine transformation events (six cassette exchanges, three in each orientation (A and B), and three transgene integrations) were detected, with a minimum cassette-exchange efficiency of 5% (6/124 F_0_) and integration efficiency of 2% (3/124 F_0_). For comparative functional analysis, representative *Cyp6* lines in orientation A were maintained and crossed with alternative GAL4 driver lines.

### CYP6M2 or CYP6P3 overexpression in multiple tissues causes distinct profiles of resistance to pyrethroids and bendiocarb

We previously described the production of a GAL4 driver line, Ubi-A10, directing widespread tissue expression (23). To quantify the overexpression achieved with this driver, we performed RT-qPCR in the progeny of Ubi-A10 driver and UAS-*Cyp6* crosses. This revealed significant 2447x (*P*=0.005) and 513x (*P*<0.001) increases of *Cyp6m2* and *Cyp6p3* transcript abundance in adult females compared to native expression in respective controls (Fig. 1A). Western analysis also readily detected CYP6M2 in the adult female progeny of the Ubi-A10/UAS-m2 crosses, but was beyond the level of detection in sibling controls (Ubi-A10/+ and +/UAS-m2) (Fig. 1B). No suitable antiserum was available for analysis of CYP6P3.

**Fig. 1.**
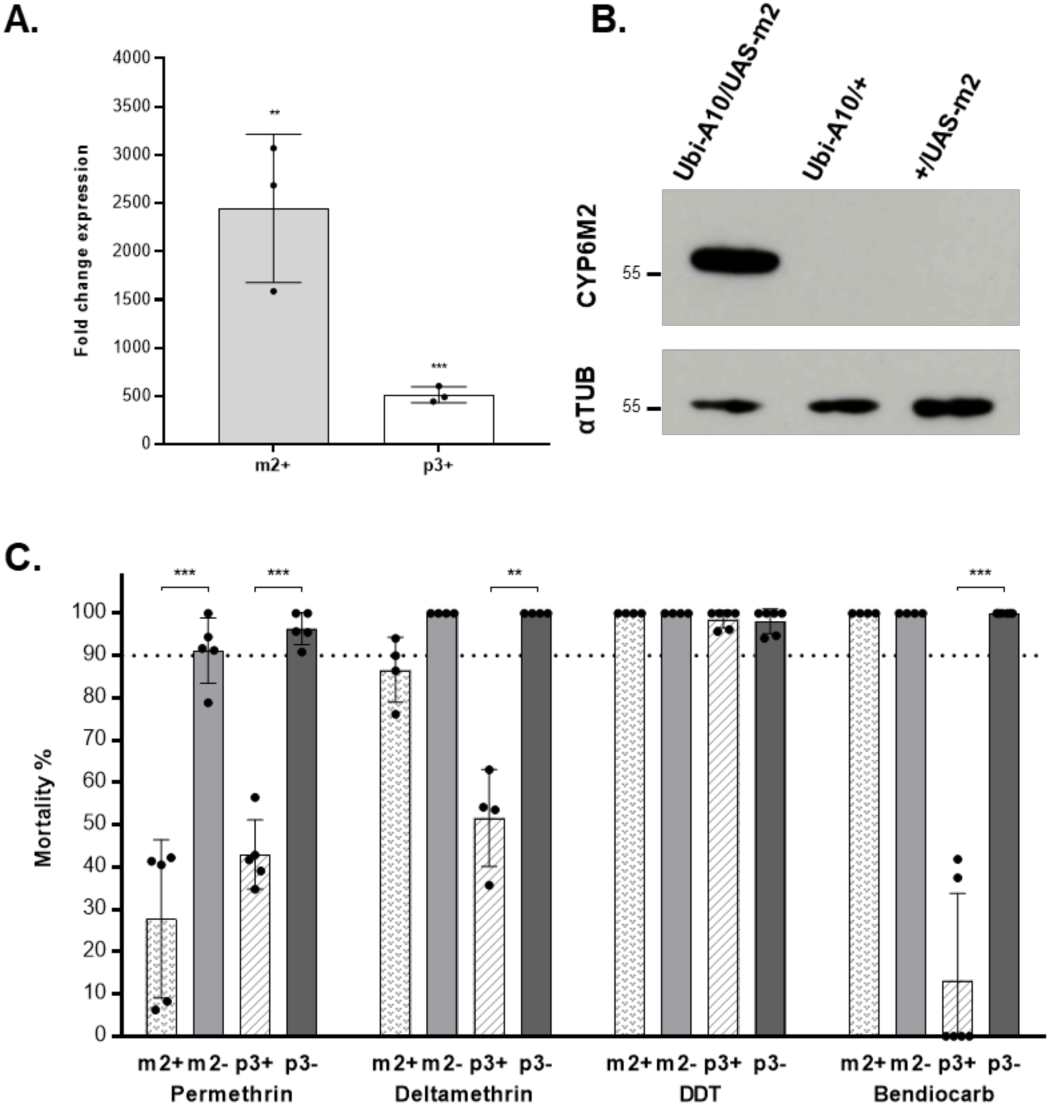
Multi-tissue *Cyp6* gene upregulation affects sensitivity to two pyrethroids and a carbamate insecticide. **A)** Relative transcription levels of *Cyp6m2* (m2+) and *Cyp6p3* (p3+) in adult females where expression is driven by the Ubi-A10 driver compared to GAL4/+ controls. Bars represent SD (N = 3). Unpaired t test, * *P*<0.05. ** *P*<0.01. *** *P*<0.001. **B)** Expression of CYP6M2 and α-tubulin in adult females from Ubi-A10 × UAS-m2 crosses with respective Ubi-A10/+ and +/UAS-m2 controls. Protein extract from the equivalent of 1/10 of a whole female mosquito was loaded in each lane. **C)** Sensitivity to insecticides of GAL4/UAS (+) females overexpressing *Cyp6m2* or *Cyp6p3* ubiquitously under the control of the Ubi-A10 driver compared to GAL4/+ controls (-) measured by WHO tube bioassay. Bars represent SD (N = 4-6, Table S2). Dotted line marks the WHO 90% mortality threshold for defining resistance. Welch’s t test with *P*<0.01 significance cut off, ** *P*<0.01, *** *P*<0.001.

WHO discriminating dose assays were then performed to assess the susceptibility of mosquitoes overexpressing *Cyp6m2* or *Cyp6p3* compared to their Ubi-A10/+ siblings. WHO tube bioassays are used to screen for the emergence of resistance in field populations and involve exposing mosquitoes to fixed concentration of insecticides (twice the LC_99_ for a susceptible strain) for 60 minutes, followed by a twenty-four-hour recovery period before recording mortality (25). The parental strains used here are susceptible (>90% mortality) to all the insecticides tested, therefore a decrease in mortality in test assays can be directly attributable to the overexpression of the specific candidate gene.

Mosquitoes overexpressing either *Cyp6* gene under the Ubi-A10 driver showed resistance to permethrin (*Cyp6m2* 28% mortality, *P*<0.001; *Cyp6p3* 43% mortality, *P*<0.001) and deltamethrin (*Cyp6m2* 88%, *P=*0.04; *Cyp6p3* 52%, *P=*0.004) compared to controls (Fig. 1C). A significant difference in mortality was observed between mosquitoes overexpressing the two different *Cyp6* genes for deltamethrin assays (*P=*0.003), while no significant difference was observed for permethrin (*P*=0.15). However, only *Cyp6p3* overexpressing mosquitoes showed resistance to bendiocarb (13% mortality *P*<0.001) (Fig. 1C). No resistance to DDT was observed with either gene in conjunction with the Ubi-A10 driver (Fig. 1C).

### CYP6M2 or CYP6P3 multi-tissue overexpression increases susceptibility to malathion

Malathion is an organophosphate pro-insecticide that is activated to a more toxic compound *in vivo* through P450-based oxidative reactions (28). Preliminary analysis at a standard WHO diagnostic dose and 60-minute exposure killed all test and control mosquitoes, however during exposure it was clear that Ubi-A10-directed *Cyp6* overexpression induced more rapid knock-down compared to controls suggesting malathion activation by these P450s. We therefore examined the relative sensitivity of mosquitoes overexpressing *Cyp6m2* or *Cyp6p3* when exposed to the same diagnostic dose of this organophosphate for a shorter time (25 minutes) (Fig. 2). Under these conditions, mosquitoes overexpressing *Cyp6m2* under the control of the Ubi-A10 driver showed significantly higher mortality rates compared to controls (95% vs 15%, *P*<0.001) and Ubi-A10/UAS-p3 mosquitoes (95% vs 34% *P*=0.002). Although, the latter also showed a trend of increased mortality compared to Ubi-A10/+ controls (34% vs 8% *P=*0.05).

**Fig. 2.**
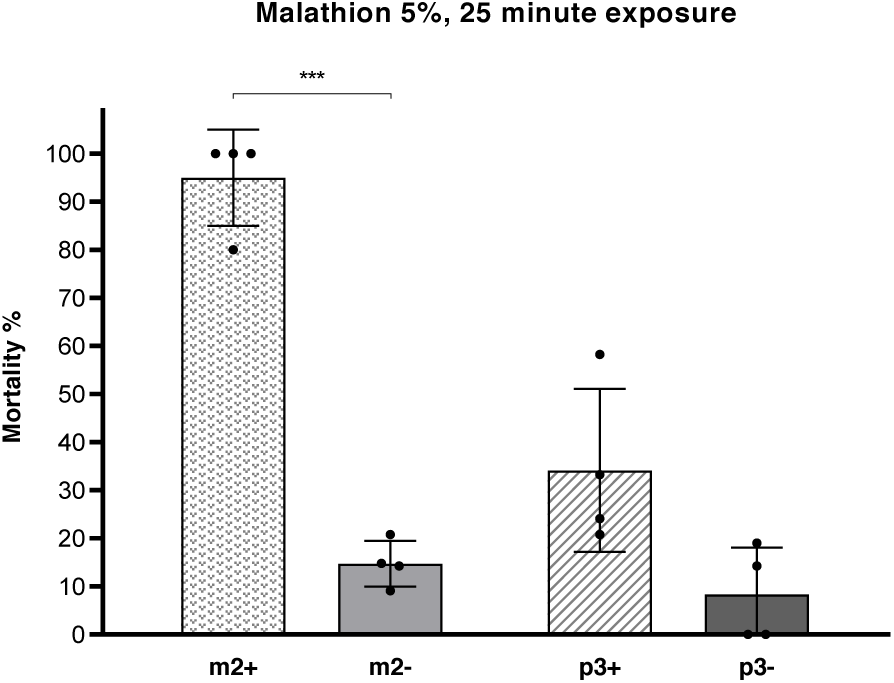
Multi-tissue *Cyp6* gene upregulation increases sensitivity to the organophosphate insecticide malathion (reduced exposure). Sensitivity to malathion of females overexpressing *Cyp6m2* (m2+) or *Cyp6p3* (p3+) ubiquitously under the control of the Ubi-A10 driver compared to respective GAL4/+ controls (m2-, p3-) measure by a modified WHO tube bioassay representing mortality rates after 25 minutes of exposure and 24 h recovery. Bars represent SD (N = 4, Table S2). Welch’s t test with *P*<0.01 significance cut off, *** *P*<0.001.

### Overexpression of GSTE2 in multiple tissues causes resistance to diagnostic doses of DDT and Fenitrothion

To extend the analysis to the role of GSTE2 in insecticide resistance in *An. gambiae*, we utilised the previously described Ubi-A10 GAL4 line (23) as a docking line for the first time. Integration of the UAS cassette into a single docking site in this case would provide Ubi-A10GAL4 and UAS-*Gste2* at the same locus (Ubi-A10GAL4:UAS-e2) and should natively overexpress *Gste2* without the need for crossing separate lines. Alternatively, cassette exchange would generate a regular UAS-*Gste2* responder line. After embryonic injections and screening, three exchange events, two in orientation A and one in orientation B (Fig. S1), and three integration events were independently recovered with an overall transformation efficiency of 9% (6/65 F_0_), exchange efficiency of 5% (3/65 F_0_), and integration efficiency of 5% (3/65 F_0_) (Table 1).

To obtain comparable data for *Gste2* and the *Cyp6* genes, we focused our analysis on the progeny from crosses between UAS-e2 and Ubi-A10GAL4 mosquitoes. When exposed to diagnostic doses of DDT, GSTE2 overexpressing mosquitoes showed a significantly lower mortality (7%, *P*<0.001) compared to controls, while no significant difference in resistance was found when exposed to diagnostic doses of permethrin, deltamethrin, malathion or bendiocarb (Fig. 3). A trend of increased tolerance was observed in mosquitoes overexpressing *Gste2* against malathion (Fig. 3), and further analysis with the related organophosphate fenitrothion indicated high resistance in Ubi-A10/UAS-e2 mosquitoes, showing 8% (*P*<0.001) mortality (Fig. 3).

**Fig. 3.**
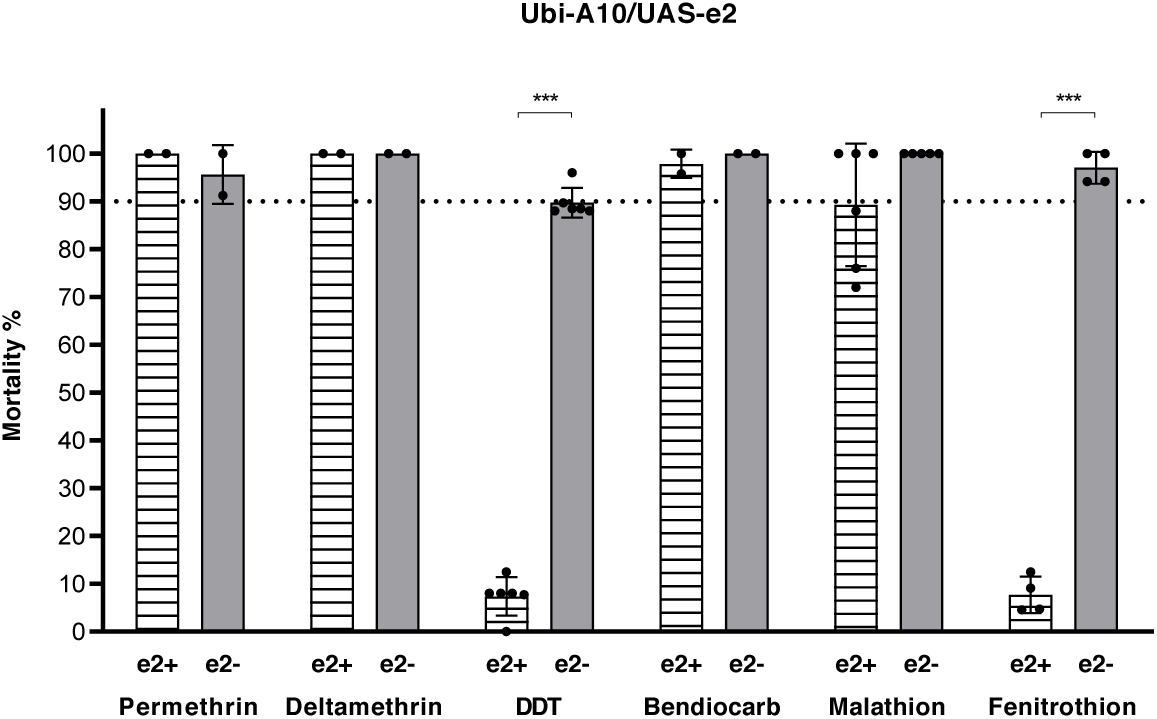
Multi-tissue overexpression of GSTE2 affects sensitivity to an organochlorine and an organophosphate insecticide. Sensitivity to insecticides of adult female mosquitoes overexpressing *Gste2* (e2+) ubiquitously under the control of the Ubi-A10 driver compared to Ubi-A10 controls (e2-) measured by WHO tube bioassay. Bars represent SD (N = 2-6, Table S2). Dotted line marks the WHO 90% mortality threshold for defining resistance. Welch’s t test with *P*<0.01 significance cut off, *** *P*<0.001.

Preliminary analysis of Ubi-A10GAL4:UAS-e2 (integration) mosquitoes indicated the expected increase in GSTE2 protein in whole body extracts compared with Ubi-A10 controls (Fig. S2A) and a resistance phenotype against DDT in the F_1_ generation of transformed male and female mosquitoes (Fig. S2B).

### Oenocyte or midgut specific overexpression of CYP6M2 or CYP6P3 does not confer resistance to insecticides

To examine the role of oenocytes and midgut tissues in P450-based metabolism of insecticides we utilised previously published GAL4 driver lines to regulate tissue specific expression. The specificity of these GAL4 drivers has been established following crosses with UAS regulated fluorescent gene reporter lines (22, 24), but to examine the relative increase in tissue-specific *Cyp6* gene expression, we performed RT-qPCR and western blot analysis in progeny from alternative driver and *Cyp6* responder crosses.

Using the midgut driver (GAL4-mid), *Cyp6m2* and *Cyp6p3* transcripts were 2730x (*P*=0.002) and 659x (*P*=0.011) more abundant in midguts dissected from GAL4/UAS mosquitoes compared to controls (Fig. 4A). A low level of overexpression was detected in the remaining carcass of GAL4/UAS mosquitoes compared to that of controls (*Cyp6m2:* 77x, *P*=0.038; *Cyp6p3:* 7x, *P*=0.08). In GAL4-oeno crosses, *Cyp6m2* and *Cyp6p3* were specifically upregulated in transgenic dissected abdomens (66x, *P*=0.013 for *Cyp6m2*; 153x, *P*<0.001 for *Cyp6p3*) where oenocytes are located (Fig. 4B). Background overexpression was also found in the remaining carcass of GAL4/UAS-m2 and -p3 adults compared to controls (26x, *P*<0.001; 2x, *P*<0.001 respectively). In western blot analysis, CYP6M2 antiserum again only detected the target protein in GAL4/UAS mosquitoes. CYP6M2 was found exclusively in dissected midguts (and whole mosquitoes) from the progeny of GAL4-mid crosses, but was not observed in GAL4/UAS carcasses or extracts from controls (Fig. 4C). Similarly, in GAL4-oeno crosses CYP6M2 signal was only detected in whole adult female extracts and in dissected abdomen integument, but not in the remaining carcass or control extracts (Fig. 4D).

**Fig. 4.**
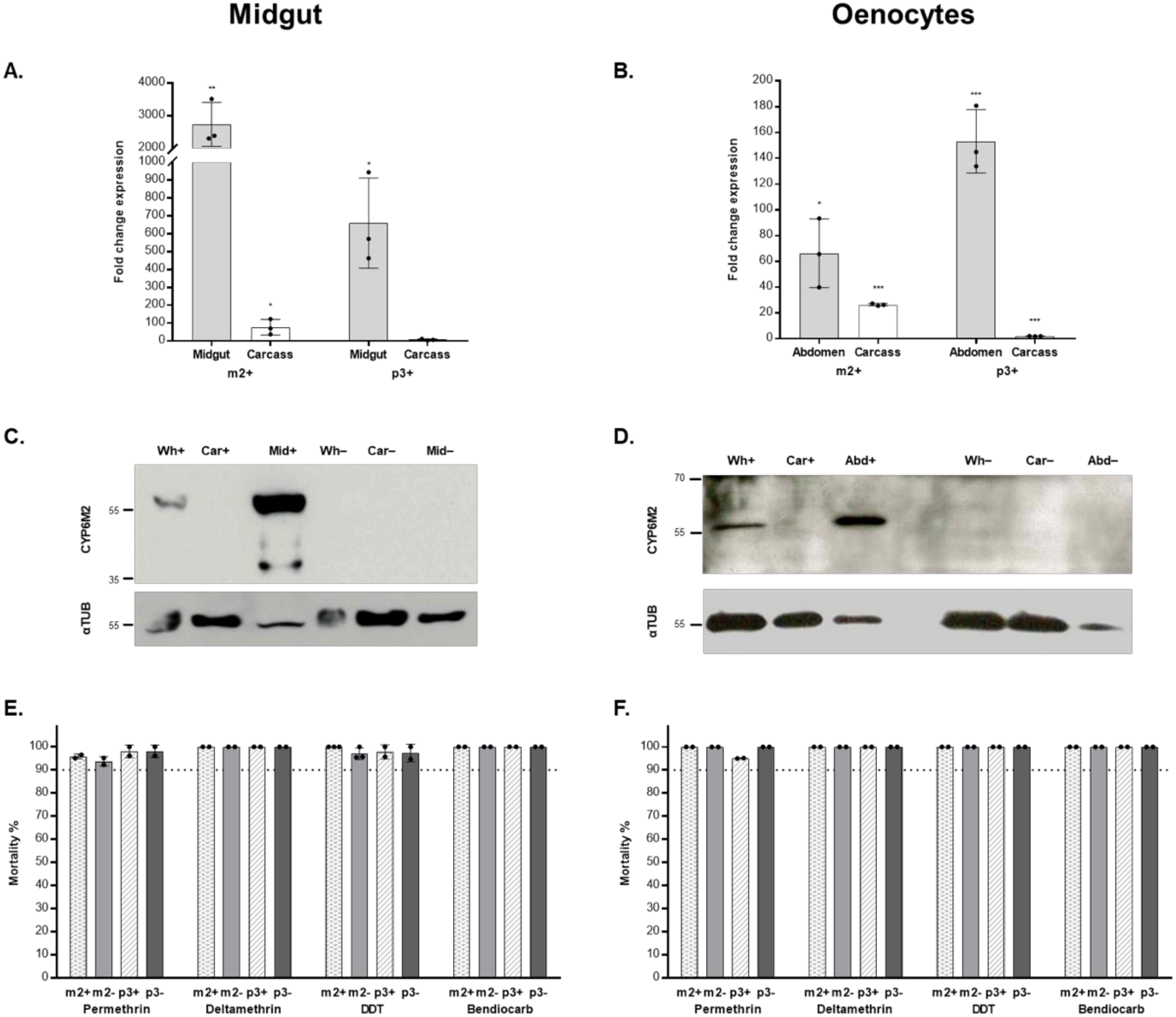
*Cyp6* gene upregulation in the mosquito midgut or oenocytes does not affect sensitivity to insecticides. **A-B)** Relative transcription levels of *Cyp6m2* (m2+) and *Cyp6p3* (p3+) in dissected midguts (A) and abdomens (B) of GAL4/UAS female mosquitoes compared to the equivalent body parts in GAL4/+ controls. Carcass is whole body without the relevant dissected part. Bars represent SD (N = 3). Unpaired t test, * *P*<0.05. ** *P*<0.01. *** *P*<0.001. **C-D)** Expression of CYP6M2 and α-tubulin in females from the GAL4-mid × UAS-m2 (C) and GAL4-oeno × UAS-m2 (D) crosses. Wh: protein extract from 1/3 of a single whole female; Car: protein extract from 1/3 of a single female carcass (whole body without midgut); Mid: two dissected midguts; Abd: abdomen cuticle; +: GAL4/UAS-m2; –: GAL4/+. **E-F)** Sensitivity to insecticides of GAL4/UAS females overexpressing (+) *Cyp6m2* or *Cyp6p3* in the midgut (E) or in the oenocytes (F) compared to GAL4/+ controls (-) measured by WHO tube bioassay. Bars represent SD (N = 2-3, Table S2). Dotted line marks the WHO 90% mortality threshold for defining resistance. Welch’s t test with *P*<0.01 significance cut off.

Adult females overexpressing *Cyp6m2* in the midgut (Fig. 4E) or in the oenocytes (Fig. 4F) showed complete susceptibility to permethrin, deltamethrin, DDT, and bendiocarb. Similar results were obtained with *Cyp6p3* (Fig. 4E and F), however potential resistance (95% mortality, *P*=0.013) was suggested in oenocyte specific *Cyp6p3* overexpressing mosquitoes when exposed to permethrin (Fig. 4F). Further analysis was performed to detect subtle differences in susceptibility by repeating the assays with reduced exposure time (Fig. S3). However, no significant decrease (*P*<0.01) was found in the mortality rates of mosquitoes overexpressing *Cyp6m2* or *Cyp6p3* in the midgut or oenocytes compared to their respective controls when exposed for 20 minutes to the same diagnostic doses of the four insecticides (Fig. S3).

Finally, the 25-minute reduced exposure bioassay for malathion showed no significant difference in the mortality of mosquitoes overexpressing *Cyp6m2* or *Cyp6p3* in midgut or oenocytes compared to controls (Fig. S4).

## DISCUSSION

*In vivo* functional analysis is critical to provide evidence of causative links between candidate genes and their proposed phenotypes. Here we demonstrate the utility of new GAL4/UAS-based tools to characterise gene function directly in *An. gambiae* by reporting the first use of the system to validate the ability of single candidate genes to confer WHO-defined resistance to different classes of insecticides. Overall, the transgenic analysis in *An. gambiae* is more in accordance with data generated from recombinant protein studies of insecticide metabolism rather than those obtained from *Drosophila* survival assays (Table 2).

**Table 2.** *In vitro* (metabolism and/or depletion) and *in vivo* (*An. gambiae and D. melanogaster*) functional validation of *An. gambiae Cyp6m2, Cyp6p3*, and *Gste2* genes.

| Class | Insecticide | Gene | *In vitro* | *An. gambiae*<br>(this study) | *Drosophila* |
| --- | --- | --- | --- | --- | --- |
| Pyrethroids | Permethrin | *Cyp6m2* | ✓ ‡ (11), § (19) | ✓ | ✓ (15) |
|  |  | *Cyp6p3* | ✓ ‡ (10), § (19) | ✓ | ✓ (15) |
|  |  | *Gste2* | N/A | ✗ | ✗ (20) |
|  | Deltamethrin | *Cyp6m2* | ✓ ‡ (11), § (19) | ✓ | ✓ (15) |
|  |  | *Cyp6p3* | ✓ ‡ (10), § (19) | ✓ | ✓ (15) |
|  |  | *Gste2* | N/A | ✗ | N/A |
| Organochlorines | DDT | *Cyp6m2* | ✗ § (19) ✓ ‡* (12) | ✗ | ✓ (15) |
|  |  | *Cyp6p3* | ✗ § (19) | ✗ | N/A |
|  |  | *Gste2* | ✓ ‡ (9, 13) | ✓ | ✓ (13, 20) |
| Carbamates | Bendiocarb | *Cyp6m2* | ✗ § (15, 19) | ✗ | ✓ (15) |
|  |  | *Cyp6p3* | ✓ § (15, 19) | ✓ | ✓ (15) |
|  |  | *Gste2* | N/A | ✗ | N/A |
| Organophosphates | Malathion | *Cyp6m2* | ✓ ‡ (28), § (19) | ✓ | N/A |
|  |  | *Cyp6p3* | ✓ § (19) | ✓ | N/A |
|  |  | *Gste2* | N/A | ✗ | N/A |
|  | Fenitrothion | *Cyp6m2* | ✓ § (19) | N/A | N/A |
|  |  | *Cyp6p3* | ✓ § (19) | N/A | N/A |
|  |  | *Gste2* | N/A | ✓ | N/A |

Presence (✓) or absence (✗) of *in vitro* activity or *in vivo* WHO-defined insecticide resistance (*An. gambiae*) or increased insecticide tolerance (*Drosophila*).
‡ metabolism; § depletion
\*in presence of added cholate

In *Anopheles*, multi-tissue overexpression of *Cyp6m2* and *Cyp6p3* demonstrated that resistance to permethrin and deltamethrin (type I and II pyrethroids respectively) can be conferred by the sole overexpression of either *Cyp6* gene. *Cyp6p3* expression also conferred resistance to bendiocarb (carbamate); while the overexpression of either *Cyp6* gene did not alter DDT (organochlorine) sensitivity. These phenotypes correlate with the profile of metabolism or substrate depletion of the respective insecticides for the two recombinant P450 enzymes (Table 2). More variable results have been observed using *Drosophila* as an *in vivo* model, with overexpression of *Cyp6m2* surprisingly generating increased tolerance to bendiocarb compared with *Cyp6p3*, despite *in vitro* analysis not detecting activity against bendiocarb for *Cyp6m2* (15, 19). DDT tolerance was also observed in *Cyp6m2* overexpressing fruitflies, but data for *Cyp6p3* could not be generated (15) (Table 2). In this study, DDT resistance was monitored by dose response assays over a 24 hr exposure time, whilst bendiocarb resistance was not observed when measured through such dose response assays but was reported following 24 hr exposure to a diagnostic dose. In the latter case, the controls used to compare *Cyp6m2* and *Cyp6p3* overexpression showed very different levels of sensitivity to bendiocarb, which appeared to contribute to the differences in resistance levels observed; whilst there was no data for the respective *Cyp6p3* controls in the DDT analysis for comparison. It may thus be a difference in genetic background that gives rise to the discrepant results observed in *Drosophila*. However, it should also be noted that the different methods of insecticide bioassay may not yield directly comparable results to the diagnostic WHO level of resistance in mosquitoes used in this study and extensively used to assess the emergence of resistance in endemic countries. Our data in mosquitoes unequivocally indicate though that the expression of single *Cyp6* genes can confer resistance to different pyrethroids, and that *Cyp6p3* overexpression confers cross resistance to prominent representatives of at least two classes of public health insecticides.

In contrast to our *Cyp6* studies, increased *An. gambiae Gste2* (*AgGste2*) expression generates clear DDT resistance, while resistance to bendiocarb and pyrethroids was not observed. These phenotypes again validate predictions from the DDT activity observed *in vitro* for recombinant AgGSTE2 (9, 13) as well as the increased DDT tolerance (13) and lack of pyrethroid tolerance (20) observed when overexpressed in *Drosophila*. The corresponding *in vitro* data for AgGSTE2 activity against bendiocarb and pyrethroids have not been reported, and this is the first time that bendiocarb resistance has been examined *in vivo* following *Gste2* overexpression.

Although DDT tolerance was also observed in *Drosophila* overexpressing the orthologous *An. funestus Gste2* (*AfGste2*) (16, 18), conflicting results were reported about activity towards pyrethroids. For example, recombinant AfGSTE2 depleted permethrin but not deltamethrin *in vitro*, yet *Drosophila* acquired increased tolerance to both insecticides when *AfGste2* was overexpressed (16, 18). RNAi analysis in deltamethrin resistant *Ae. aegypti* of *AaGste2* has also indicated a role in pyrethroid resistance (29). It is possible that the variation observed in resistance profiling are due to intrinsic differences in the activity of GSTE2s derived from the different mosquito species. In this context, it has been speculated that the predominant pyrethroid detoxification role of GSTs in some insects is sequestration or protection against oxidative stress rather than direct metabolism (30). Our results show that even high levels of *AgGste2* overexpression do not confer WHO diagnostic levels of resistance to this class of insecticides in isolation. It is possible that *AgGste2* may need to work in concert with other genes, that are not upregulated in the sensitive genetic background of the *An. gambiae* transgenic lines, to produce a pyrethroid resistance phenotype. Future work will test this hypothesis by co-expression of other UAS regulated detoxification genes using the Ubi-A10GAL4:UAS-e2 (integration) line. Although beyond the scope of this work, this mosquito line expresses GAL4 and GSTE2 and can be crossed with other UAS lines to provide co-expression with other detoxification enzymes to examine additive or synergistic interactions.

Although GSTs have been associated with organophosphate (OP) metabolism through biochemical studies (7), we report the first evidence that the expression of a single gene can provide OP resistance in mosquitoes. The high resistance shown towards fenitrothion by *Gste2* overexpressing *An. gambiae* is intriguing. It is currently unclear if GSTE2 detoxifies fenitrothion by sequestration, free radical protection or directly through conjugation/modification. Evidence from early studies (31) suggest that *Anopheles* GST activity is associated with the conversion of fenitrothion to the non-toxic metabolite desmethyl fenitrooxon through an oxidised intermediate. Similar analysis in the *Gste2* overexpressing lines would clarify which of these mechanisms is involved. Further investigation is also needed on the OP malathion, for which we report suspected resistance when *Gste2* is overexpressed.

We have also demonstrated that *Cyp6* overexpression increases susceptibility to malathion, as well as conferring permethrin resistance, which may have direct implications on insecticide management, especially if replicated with other OPs that may be used for *Anopheles* control (32). Such sensitivity profiles are readily explained by the bio-activation of malathion to its more toxic metabolite malaoxon (33) by a P450-mediated mechanism (28). Here we provide the first direct *in vivo* evidence that CYP6 enzymes can confer negative cross resistance. Furthermore, there appears to be substrate specificity in the alternative P450-mediated reactions, since we observed higher mortality when assayed against *Cyp6m2* overexpression compared to *Cyp6p3*. This may suggest that *Cyp6m2* favours the higher steady state production of the toxic intermediate compared to *Cyp6p3*.

Malathion activation by *Cyp6m2* is also supported by recent evidence provided by Ingham et al (34) who found that knock down of the transcription factor Maf-S results in increased survival following malathion exposure. One of the P450s downregulated by Maf-S knockdown was *Cyp6m2*, whereas *Cyp6p3* transcription was not modified. Taken together, the results provide experimental evidence to support the use of OPs, and potentially other pro-insecticides activated by CYP6 enzymes, for *Anopheles* control in areas where pyrethroid resistance is also conferred by detoxification by the same enzyme/s. One such strategy involves combining the use of pyrethroid-based bed nets with OP-based residual wall spraying or impregnated hangings (32). This takes advantage of the additive effect of the two classes of insecticides, while sensitising *Cyp6*-based pyrethroid resistant mosquitoes to malathion (35). In conjunction with recombinant enzyme assays, the modified mosquitoes described may thus become valuable tools to assess the susceptibility of new public health pro-insecticides, for example chlorfenapyr (36), to activation and detoxification by xenobiotic metabolising P450 genes in *Anopheles*.

When validating resistance phenotypes conferred by transgenic overexpression, the spatial pattern of overexpression can give clues to the identity of key tissues of detoxification. The expression driven by Ubi-A10 is spread over multiple tissues, which makes it impossible to pinpoint which tissue/s are particularly important for generating the resistance phenotype. Here, we directly investigated the involvement of the midgut and oenocytes in conferring P450-mediated resistance. Critically, we did not observe clear resistance to any insecticide class when either *Cyp6m2* or *Cyp6p3* were specifically expressed in either of these tissues, despite achieving highly enriched expression and the knowledge that oenocytes and the midgut express abundant P450 co-enzyme CPR (26). Furthermore, since our previous expression profiling of the Ubi-A10 driver indicated lack of expression in Malpighian tubules (23) yet resistance to multiple insecticides was observed with this driver, it would appear that the insecticides tested are not predominately metabolised in the Malpighian tubules either, and other unidentified tissues may be critical, alone or in combination, for detoxification. As described earlier, some evidence of tissue specificity of P450s associated with insecticide resistance has been derived from transcriptomic analysis of crude dissections of tissues and body segments from pyrethroid resistant and sensitive strains (27). This study indicated that *Cyp6p3* is more highly expressed in the midgut of the resistant strain, whereas *Cyp6m2* has a broader upregulation in midgut, Malpighian tubules and the abdomen (integument, fat body and ovaries). The relevance of elevated *Cyp6p3* levels in the midgut of the examined resistant strain is difficult to reconcile with the lack of a resistance phenotype when the same gene is overexpressed in this tissue with the GAL4/UAS system.

Previous *Drosophila* studies have shown that overexpression using drivers active in multiple tissues, such as actin5C-GAL4 (14–18) or tubulin-GAL4 (20) are generally needed to modify resistance. Nevertheless, there are few examples in which tissue-specific drivers have been used to validate *Cyp6* gene based resistance in *Drosophila*. Yang et al (37) demonstrated the central role of Malpighian tubules for *DmCyp6g1*-mediated DDT resistance, whilst Zhu et al (38) demonstrated the importance of neuronal expression to provide deltamethrin resistance in *Drosophila* expressing *T. castaneum Cyp6bq9*. Even in this latter analysis, however, the neuronal driver showed leaky expression in other tissues, leading to the possibility that the observed phenotype results from expression in multiple tissues. Overall, a more detailed analysis with further tissue-specific drivers, as they become available, is needed to clarify the potential involvement of specific tissues in the detoxification of insecticides in *An. gambiae*.

## Conclusions

This work reports on the first functional analysis of mosquito insecticide resistance genes conducted in transgenic *An. gambiae*. The mosquitoes generated are resistant, in a solely metabolism-based manner, to at least one representative insecticide from the major classes used in public health, and are therefore useful in liability screens of new and repurposed active compounds, including insecticides, pro-insecticides, synergists and sterilising agents. The lines can also be used in combination with strains carrying genome edited target sites (e.g. Kdr and Ace-1R) to examine the additive or synergistic effects of multiple resistance mechanisms. Similarly, it is possible to use the integration line carrying both Ubi-GAL4 and UAS-*Gst*e2 to cross with other UAS-detox genes to analyse metabolic interactions, for example combining phase I and II metabolism. In addition, the Ubi-A10 driver is active in larval stages (23) and can thus be used to examine gene function in immature stages.

Importantly, for future work, there is growing evidence on the involvement in resistance of genes that are very difficult to test *in vitro* due to the lack of appropriate assays. These include genes coding for cuticle components (39), transcription factors (34) and other binding proteins, e.g. hexamerins and α-crystallins (21), for which current transgenic tools, including GAL4/UAS, make *An. gambiae* the most relevant option for functional genetic analysis.

## MATERIALS AND METHODS

### Plasmid construction

Responder plasmids were designed for the expression of the *An. gambiae* genes *Cyp6m2* (AGAP008212), *Cyp6p3* (AGAP002865), or *Gste2* (AGAP009194) under the regulation of the UAS and carried a YFP marker gene regulated by the 3xP3 promoter. The coding sequences of *Cyp6m2* (1500 bp), derived from the susceptible strain Kisumu, was amplified from PB13:CYP6M2 (11) using primers M2fw and M2rv (Table S1). The coding sequence of *Cyp6p3* was obtained by amplifying a 193 bp fragment from Kisumu cDNA using primers P3fw1 and P3rv1 (Table S1) and a 1362 bp fragment from pCW:17α-*Cyp6p3* (10) using primers P3fw2 and P3rv2 (Table S1). P3fw1 and P3rv2 were then used to join the two fragments and obtain the 1530 bp full length *Cyp6p3* coding sequence. The 666 bp *Gste2-114T* coding sequence derived from the DDT-resistant strain ZAN/U was amplified from the K1B plasmid (13) using primers Gste2k1bfor and Gste2k1brev (Table S1). All coding sequences were cloned into the YFP-marked responder plasmid pSL*attB:YFP:Gyp:UAS14i:Gyp:attB (24) downstream of the UAS using EcoRV/XhoI (*Cyp6*) or EcoRI/NcoI (*Gste2*).

### Creation of UAS responder lines by PhiC31-mediated cassette exchange

For creating responder lines carrying *Cyp6* genes, embryos of the docking line A11 (24), which carries two inverted *attP* sites and is marked with 3xP3-driven CFP, were microinjected with 350 ng/µl of the responder plasmid and 150 ng/µl of the integrase helper plasmid pKC40 encoding the phiC31 integrase (40) as described in Pondeville et al (41). The same protocol was followed to create the *Gste2* responder line using embryos of the docking line Ubi-A10 (23) which carries two inverted *attP* sites and is marked with 3xP3-driven CFP. Emerging F_0_ were pooled into sex specific founder cages and outcrossed with wild type G3s. F_1_ progenies were screened for the expression of YFP (cassette exchange) and CFP/YFP (cassette integration) in the eyes and nerve cord. Orientation check to assess the direction of cassette exchange was performed on F_1_ YFP-positive individuals or on the F_2_ progeny deriving from single YFP-positive individuals. This was carried out by PCR using alternative combinations of four primers designed to give a product only in one of the orientations: PiggyBacR-R2 + Red-seq4R (PCR1) and M2intFW or P3intFW or Gste2_v1 + ITRL1R (PCR2) to detect insertions in orientation A; PiggyBacR-R2 + M2intFW or P3intFW or Gste2_v2 (PCR3) and Red-seq4R + ITRL1R (PCR4) for orientation B. All definitive responder lines were created from individuals showing orientation of insertion A, which was chosen for consistency with previous RMCE lines created in this laboratory. Transformation efficiencies were calculated as the number of independent transgenic events (exchanges or integrations) over the number of surviving F_0_ adults.

### Driver lines and GAL4 x UAS crosses

Crosses for ubiquitous expression were established between the CFP-marked driver Ubi-A10 (23) and individuals of the responder lines marked with YFP. While to obtain tissue-localised expression dsRed-marked drivers specific for expression in the midgut (GAL4-mid) (22) or in the oenocytes (GAL4-oeno) (24) were used. Responder lines were kept as a mix of homozygous and heterozygous individuals so to obtain GAL4/+ progeny to be used as transgenic blank controls.

### Cyp6 gene expression analysis

To quantify *Cyp6* gene expression in GAL4/UAS and GAL4/+ individuals, total RNA was harvested from pools of 2-5-day-old whole adults and their relevant dissected body part (midgut or abdomen cuticle). The adult tissues remaining after dissection constituted the carcass. Three biological replicates consisting of 5 mosquitoes (or body parts) each were collected from each mosquito population. RNA extraction was performed using the TRI Reagent^®^ protocol (Sigma). To remove genomic DNA contamination, samples were treated with the Turbo DNA-Free kit (Ambion). RNA was then reverse-transcribed using the SuperScript III First-Strand Synthesis System (Life Technologies) following the oligo(dT) reaction protocol. RT-qPCR reactions were set up using 1x Brilliant III Ultra-Fast SYBR^®^ Green qPCR Master Mix (Agilent Technologies) and primers qM2fw and qM2rv for quantification of *Cyp6m2*, and qP3fw and qP3sub for *Cyp6p3* (15) (Table S1). The qP3sub primer bears a nucleotide substitution (A11G) to conform its sequence to that of the G3 strain template. Two housekeeping genes, the ribosomal protein S7 (RPS7) (AGAP010592) and ribosomal protein L40/Ubiquitin (AGAP007927), were also quantified using primers qS7fw, qS7rv, qUBfw and qUBrv (42) (Table S1). Transcription data obtained by RT-qPCR were analysed using the ΔΔCt method as described in SI. Gene expression analysis was not performed to assess upregulation of the *Gste2* transcript.

### CYP6 and GSTE2 protein expression analysis

To detect protein expression in GAL4/UAS and GAL4/+ individuals, total protein extracts were obtained from whole 2-5-day-old female adults and their dissected body parts. Protein extracts equivalent to 1/3 of a mosquito or its body part were analysed to detect CYP6 expression driven by tissue-specific drivers. With the exception of midgut samples, for which two whole midguts were analysed. The higher amount of midgut sample was required to visualise signal of the α-tubulin loading control. The equivalent of 1/10 of a single female mosquito was used to assess expression driven by ubiquitous drivers. CYP6s were probed using primary affinity-purified polyclonal peptide antibodies produced in rabbit against CYP6M2 or CYP6P3 (gifts from Dr M. Paine), while GSTE2 was probed with anti-GSTE2-28 rabbit primary antibodies (9). Secondary antibodies were anti-rabbit-HRP IgGs (Bethyl Laboratories). Detection of the loading control α-tubulin was performed using primary mouse anti-αtubulin antibodies (Sigma or Fisher Scientific) and secondary goat anti-mouse-HRP IgG antibodies (Abcam). Signal detection was carried out using SuperSignal™ West Dura Extended Duration Substrate (Life Technologies).

### Assessment of susceptibility to insecticides

Susceptibility to insecticides was assessed in mosquitoes overexpressing *Cyp6* genes using the WHO tube bioassay (25). Pools of 20-25 GAL4/UAS and GAL4/+ adult female mosquitoes were exposed 2-5 days post-emergence to standard discriminating doses of insecticides – 0.75% permethrin, 0.05% deltamethrin, 0.1% bendiocarb, 4% DDT – for 60 minutes and mortality rates assessed after a 24 hour recovery period. For mosquitoes expressing *Cyp6* genes in the midgut or oenocytes a modified version of the standard WHO test was also performed reducing the exposure time to 20 minutes (26). For assessing susceptibility to 5% malathion in mosquitoes overexpressing *Cyp6* genes, the exposure time was decreased to 25 minutes. Mosquitoes overexpressing *Gste2* were additionally tested for 1% fenitrothion using the recommended 2 h exposure time. 1-4 biological replicates were performed for each insecticide tested. A total of 2-8 technical replicate tubes were tested for each population. Welch’s t-test was performed to determine statistical differences between mortality rates in GAL4/UAS and GAL4/+. Details on statistical analysis and replicate numbers of bioassay experiments are reported in Table S2.

## ACKNOWLEDGMENTS

We gratefully acknowledge the LSTM and the MRC (MR/P016197/1) for sponsoring PhD studentships to AA and BP respectively, and to the Innovative Vector Control Consortium for follow on funding to AA. Thanks also go to Dave Weetman for very useful comments on the manuscript.

