## supplementary information for "Functional genetic validation of key genes conferring insecticide resistance in the major African malaria vector, *Anopheles gambiae*"

**This PDF file includes:**

Supplementary text: *Cyp6* expression analysis using the ΔΔCt method

Figs. S1 to S4

Tables S1 and S2

### Supplementary Text

#### *Cyp6* expression analysis using the ΔΔCt method

Threshold cycle (Ct) values were corrected for primer efficiency applying the formula *Corrected Ct = Ct (Logx / Log2)*, where x = 2 if efficiency is >100% and x = 1+Efficiency if efficiency is <100%.

Ct values obtained for target genes were normalised against those of the housekeeping genes RPS7 and Ubiquitin using the formula *ΔCt = Ct target – Ct housekeeping.*

Mean ΔCt values of GAL4/+ mosquitoes were then compared to those of GAL4/UAS mosquitoes using the formula *ΔΔCt = ΔCt GAL4/+ – ΔCt GAL4/UAS*. Fold change (FC, x) expression between the two populations was calculated with the formula *FC = (2^ GAL4/UAS ΔΔCt) / (2^ GAL4/+ ΔΔCt)*.

Statistical differences in fold change expression between GAL4/UAS and GAL4/+ were determined using t test (unpaired, two-tails, assuming equal variance).

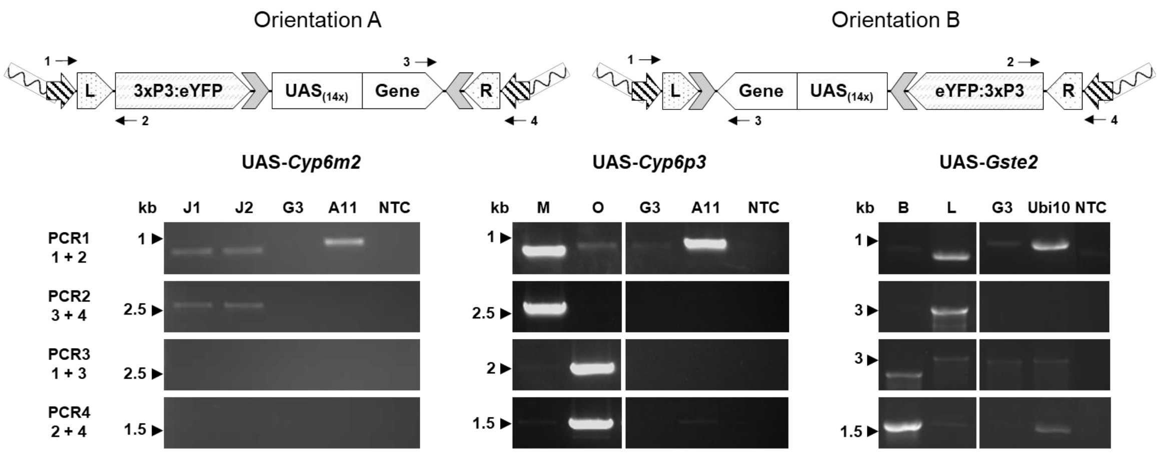


**Fig. S1. Orientation of insertion of the UAS transgenic cassette in representative individuals of responder lines.** Two orientations of insertion, A and B, are possible after recombination of the *attP* sites in the docking mosquito genome and the attB sites in the UAS plasmids. Striped arrows: piggyBac transposon arms; 3xP3: eye and nerve-cord specific promoter; eYFP: enhanced yellow fluorescent protein; grey arrows: Gypsy insulators; $UAS_{(14x)}$: upstream activating sequence (14 repeats). Each PCR (1-4) gives a distinct amplification fragment that is diagnostic for the orientation of the insertion. →: primer annealing site. 1: PiggyBacR-R2, 2: Red-seq4R, 3: M2intFW or P3intFW or GSTe2_v1/v2, 4: ITRL1R. L, R: attL and attR hybrid sites created after recombination. J1, J2: individuals from the UAS-*Cyp6m2* line; M, O: representative individuals from the UAS-*Cyp6p3* line; B, L: representative individuals from the UAS-*Gste2* line; G3: wild type; A11: docking line control; Ubi10: docking line control; NTC: negative control (water as template).

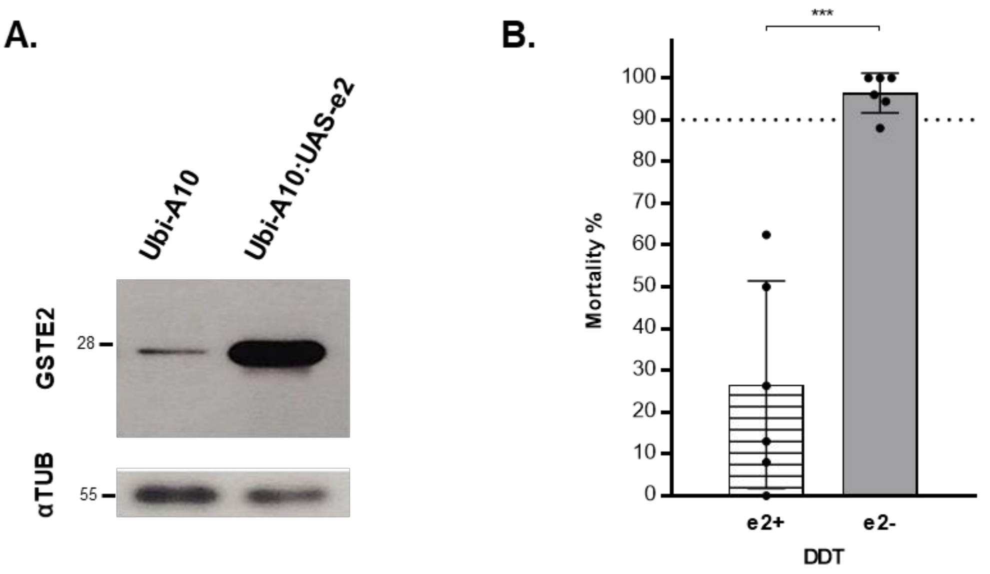


**Fig. S2. Multi-tissue overexpression of GSTE2 in the integration line affects sensitivity to the organochlorine insecticide DDT. A)** Expression of GSTE2 and α-tubulin in adult females of the Ubi-A10:UAS-e2 line with respective Ubi-A10/+ controls. Protein extract from the equivalent of 1/10 of a whole female mosquito was loaded in each lane. **B)** Sensitivity to DDT of Ubi-A10:UAS-e2 females overexpressing *Gste2* (e2+) ubiquitously under the control of the Ubi-A10 driver compared to Ubi-A10/+ controls (e2-) measured by WHO tube bioassay. Bars represent SD (N = 6, Table S2). Dotted line marks the WHO 90% mortality threshold for defining resistance. Welch's t test with $P<0.01$ significance cut off, *** $P<0.001$.

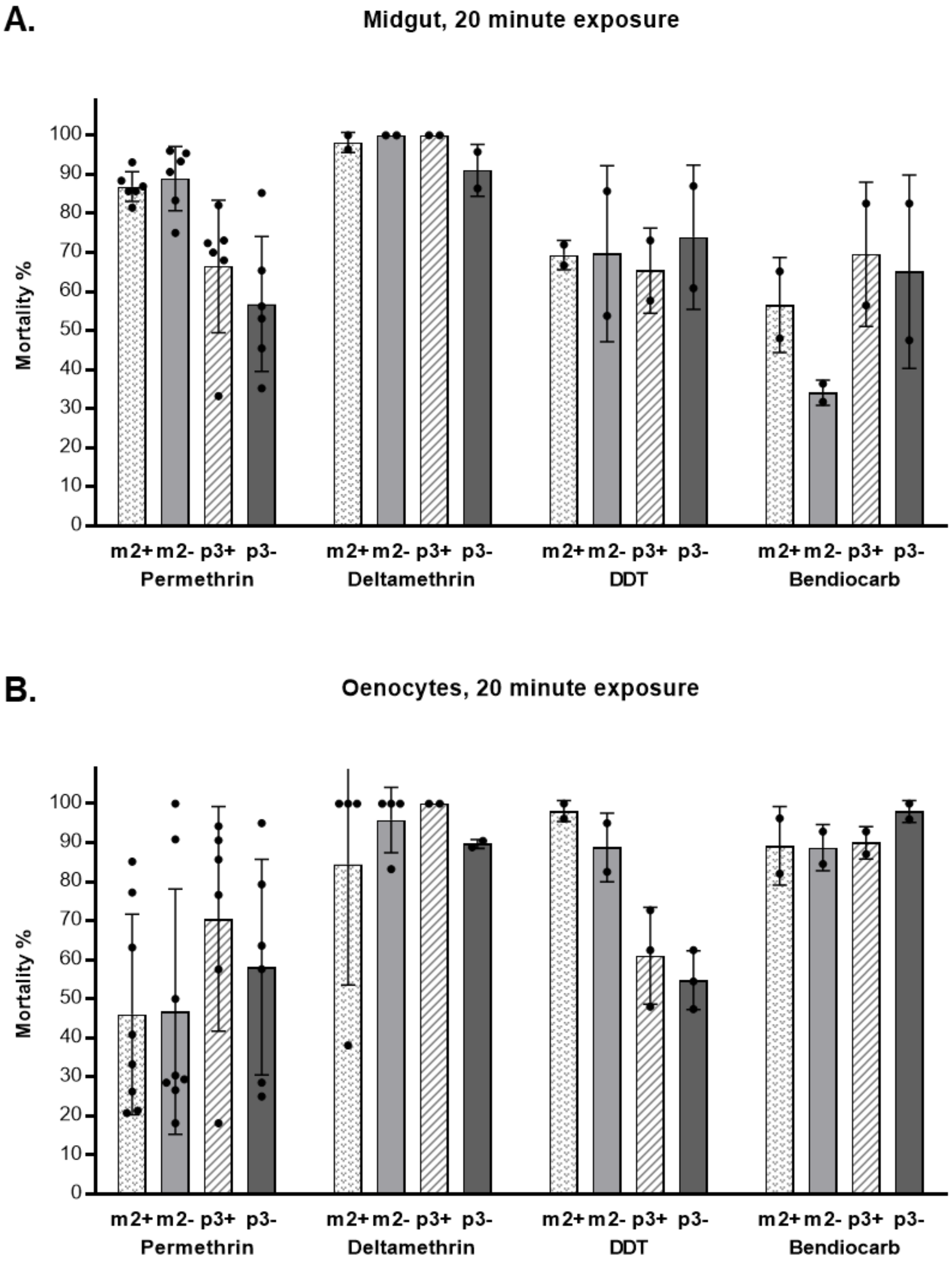


677 **Fig. S3. Tissue-specific *Cyp6* gene upregulation does not affect sensitivity to insecticides**
678 **(reduced exposure).** Sensitivity to insecticides of GAL4/UAS (+) females overexpressing
679 *Cyp6m2* or *Cyp6p3* under the control of the midgut (A) or oenocyte specific (B) drivers compared
680 to GAL4/+ controls (-) measured by a modified WHO tube bioassay representing mortality rates
681 after 20 minutes of exposure and 24 h recovery. Bars represent SD (N = 2-8, Table S2). Welch's t
682 test with $P$<0.01 significance cut off.

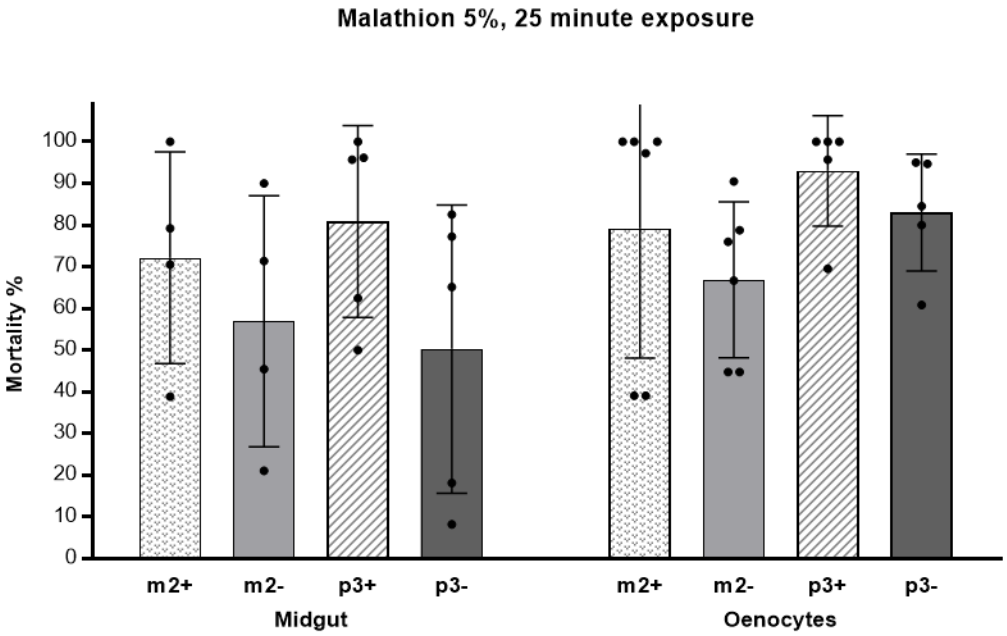


**Fig. S4. Tissue-specific *Cyp6* gene upregulation does not affect sensitivity to the organophosphate insecticide malathion (reduced exposure).** Sensitivity to malathion (reduced exposure) of females overexpressing *Cyp6m2* (m2+) or *Cyp6p3* (p3+) tissue-specifically (midgut, oenocytes) compared to respective controls (m2-, p3-) measure by modified WHO tube bioassay representing mortality rates after 25 minutes of exposure and 24 h recovery. Bars represent SD (N = 4-6, Table S2). Welch's t test with $P$<0.01 significance cut off.

**Table S1. Primers used in this study**

| Primer | Sequence 5'-3' |
|---|---|
| **M2fw** | TTCTGATATCAAAAATGTTTAGCTTGTTGGATTTCA |
| **M2rv** | TTCTCTCGAGCTAAATCTTATCCACCTTCAACCAC |
| **P3fw1** | TTCTGAATTCAACGATGGAGCTAATTAACGCGGTGCTGG |
| **P3rv1** | GGTACAGCTCCTGATGGATGTCGGC |
| **P3fw2** | GCCGACATCCATCAGGAGCTGTACC |
| **P3rv2** | TTCTCTCGAGCTACAACTTTTCCACCTTCAAG |
| **Gste2k1bfor** | GGGGGAATTCGAAAATGTCCAACCTTGTACTGTACAC |
| **Gste2k1brev** | GGGGCTCGAGTTAAGCCTTAGCATTCTCCTCCTT |
| **PiggyBacR-R2** | TTTGCCTTTCGCCTTATTTTAGA |
| **Red-seq4R** | CGAGGGTTCGAAATCGATAA |
| **M2intFW** | CGTATAGGGCTGGCGTATCT |
| **P3intFW** | GCTGAGAAAGTTCCGCTTCT |
| **GSTe2_v1** | TGTAAATTCGGCCCTGCACT |
| **GSTe2_v2** | GTGTAAATTCGGCCCTGCAC |
| **ITRL1R** | TGACGAGCTTGTTGGTGAGGATTCT |
| **qM2fw** | TACGATGACAACAAGGGCAAG |
| **qM2rv** | GCGATCGTGGAAGTACTGG |
| **qP3fw** | TGTGATTGACGAAACCCTTCGGAAG |
| **qP3sub** | ATAGTCCACAGATGGTACGCGGG |
| **qS7fw** | AGAACCAGCAGACCACCATC |
| **qS7rv** | GCTGCAAACTTCGGCTATTC |
| **qUBfw** | CGACTCCGTGGTGGTATCAT |
| **qUBrv** | GCACTTGCGGCAAATCATCT |

### Table S2. Bioassay experiments

| | Cross | Insecticide | Standard exposure | | | | | Reduced exposure | | | | |
|---|---|---|---|---|---|---|---|---|---|---|---|---|
| | | | Experiment (Biol. reps) | Tot. tech. reps | *P* | t | df | Experiment (Biol. reps) | Tot. tech. reps | *P* | t | df |
| UBIQUITOUS | Ubi-A10 x UAS-m2 | Permethrin 0.75% | 2 | 5 | 0.0007 | 7 | 5.3 | NS | NS | NS | NS | NS |
| | | Deltamethrin 0.05% | 2 | 4 | 0.04 | 3.5 | 3 | NS | NS | NS | NS | NS |
| | | DDT 4% | 1 | 4 | N/A | N/A | N/A | NS | NS | NS | NS | NS |
| | | Bendiocarb 0.1% | 1 | 4 | N/A | N/A | N/A | NS | NS | NS | NS | NS |
| | | Malathion 5% | NS | NS | NS | NS | NS | 2 | 4 | <0.0001 | 14.5 | 4.3 |
| | Ubi-A10 x UAS-p3 | Permethrin 0.75% | 3 | 5 | <0.0001 | 13.3 | 5.6 | NS | NS | NS | NS | NS |
| | | Deltamethrin 0.05% | 1 | 4 | 0.004 | 8.4 | 3 | NS | NS | NS | NS | NS |
| | | DDT 4% | 2 | 6 | 0.7 | 0.3 | 9.1 | NS | NS | NS | NS | NS |
| | | Bendiocarb 0.1% | 2 | 6 | 0.0001 | 10.3 | 5 | NS | NS | NS | NS | NS |
| | | Malathion 5% | NS | NS | NS | NS | NS | 2 | 4 | 0.05 | 2.6 | 4.8 |
| | Ubi-A10 x UAS-e2 | Permethrin 0.75% | 1 | 2 | 0.5 | 1 | 1 | NS | NS | NS | NS | NS |
| | | Deltamethrin 0.05% | 1 | 2 | N/A | N/A | N/A | NS | NS | NS | NS | NS |
| | | DDT 4% | 3 | 6 | <0.0001 | 39.6 | 9.4 | NS | NS | NS | NS | NS |
| | | Bendiocarb 0.1% | 1 | 2 | 0.5 | 1 | 1 | NS | NS | NS | NS | NS |
| | | Malathion 5% | 3 | 6 | 0.1 | 2. | 5 | NS | NS | NS | NS | NS |
| | | Fenitrothion 1% | 2 | 4 | <0.0001 | 35.2 | 5.9 | NS | NS | NS | NS | NS |
| | Ubi-A10:UAS-e2 | DDT 4% | 3 | 6 | 0.0008 | 6.8 | 5.4 | NS | NS | NS | NS | NS |
| MIDGUT | GAL4-mid x UAS-m2 | Permethrin 0.75% | 1 | 2 | 0.35 | 1.3 | 1.4 | 3 | 6 | 0.6 | 0.6 | 7.1 |
| | | Deltamethrin 0.05% | 1 | 2 | N/A | N/A | N/A | 1 | 2 | 0.5 | 1 | 1 |
| | | DDT 4% | 2 | 3 | 0.18 | 2 | 2 | 1 | 2 | 0.98 | 0.02 | 1.1 |
| | | Bendiocarb 0.1% | 1 | 2 | N/A | N/A | N/A | 1 | 2 | 0.2 | 2.5 | 1.1 |
| | | Malathion 5% | NS | NS | NS | NS | NS | 2 | 4 | 0.5 | 0.8 | 5.8 |
| | GAL4-mid x UAS-p3 | Permethrin 0.75% | 1 | 2 | >0.99 | 0 | 2 | 3 | 6 | 0.35 | 0.98 | 10 |
| | | Deltamethrin 0.05% | 1 | 2 | N/A | N/A | N/A | 1 | 2 | 0.3 | 1.9 | 1 |
| | | DDT 4% | 1 | 2 | 0.9 | 0.1 | 1.9 | 1 | 2 | 0.6 | 0.6 | 1.6 |
| | | Bendiocarb 0.1% | 1 | 2 | N/A | N/A | N/A | 1 | 2 | 0.9 | 0.2 | 1.8 |

| | | | | | | | | | | | | |
|---|---|---|---|---|---|---|---|---|---|---|---|---|
| | | Malathion 5% | NS | NS | NS | NS | NS | 3 | 5 | 0.1 | 1.6 | 7 |
| OENOCYTES | GAL4-oeno x UAS-m2 | Permethrin 0.75% | 1 | 2 | N/A | N/A | N/A | 4 | 8 | 0.96 | 0.05 | 13.5 |
| | | Deltamethrin 0.05% | 1 | 2 | N/A | N/A | N/A | 2 | 4 | 0.5 | 0.7 | 3.4 |
| | | DDT 4% | 1 | 2 | N/A | N/A | N/A | 1 | 2 | 0.4 | 1.4 | 1.2 |
| | | Bendiocarb 0.1% | 1 | 2 | N/A | N/A | N/A | 1 | 2 | 0.96 | 0.05 | 1.6 |
| | | Malathion 5% | NS | NS | NS | NS | NS | 3 | 6 | 0.4 | 0.8 | 8.2 |
| | GAL4-oeno x UAS-p3 | Permethrin 0.75% | 1 | 2 | 0.013 | 49 | 1 | 3 | 6 | 0.5 | 0.8 | 10 |
| | | Deltamethrin 0.05% | 1 | 2 | N/A | N/A | N/A | 1 | 2 | 0.05 | 12.9 | 1 |
| | | DDT 4% | 1 | 2 | N/A | N/A | N/A | 2 | 3 | 0.5 | 0.7 | 3.3 |
| | | Bendiocarb 0.1% | 1 | 2 | N/A | N/A | N/A | 1 | 2 | 0.2 | 2.3 | 1.8 |
| | | Malathion 5% | NS | NS | NS | NS | NS | 3 | 5 | 0.3 | 1.2 | 8 |

N/A: mortality is 100% in all replicates; NS: not screened.

Each experiment included mosquitoes deriving from independent crosses or from subsequent gonotrophic cycles of the same cross, therefore they were considered as biological replicates. 1-4 experiments were performed for each insecticide tested. Generally, two technical replicate tubes containing 20-25 females were tested for each population in each experiment. Standard exposure was 60 minutes for all insecticides except fenitrothion for which recommended exposure is 2 hours. Reduced exposure was 20 minutes for permethrin, deltamethrin, DDT and bendiocarb, and 25 minutes for malathion. Recovery time is 24 hours for all experiments.
